## Supplementary Methods and Figures for "Analysis of RyR2 distribution in HEK293 cells and mouse cardiac myocytes using 3D MINFLUX microscopy"

### Supplementary Methods and supplementary Figures

#### Supplementary Methods

##### Analysis of 3D MINFLUX data

For data analysis and visualization, raw MINFLUX localization data were exported using Abberior Instruments Imspector software[1] (MINFLUX version) to Numerical Python (Numpy) NPY format. The NPY formatted data was directly imported into PYMEVisualize[2] using custom plugin code in the `PYMEcs.IO.MINFLUX` module of the PYME-extra repository (available on github at github.com/csoeller/PYME-extra). Upon importing the data a foreshortening factor of 0.72 was applied to scale all z-coordinates, similar as described previously[3], and registered in the metadata. This factor accounts for refractive index mismatch between the coverslip glass and cell material and was determined by measuring the NPC ring spacing in 3D MINFLUX data from U2OS-Nup96-mEGFP cells to give a mean of ~50 nm after foreshortening correction.

###### *Estimating docking strand and RyR2 subunit locations*

*Raw MINFLUX localizations* were processed by combining localizations to represent different aspects of the underlying structures. During attachment to the docking strand on sdABs an emitting imager strand is typically localized several times in direct succession by rapidly repeating MINFLUX localization iterations, until the imager strand eventually unbinds. These successive localizations are assigned to the same "trace" via the same "trace ID" (exported parameter TID)[4]. Localization precision was estimated as the standard deviation of MINFLUX localization coordinates of localizations belonging to the same TID[3,4], limiting localizations to traces that contain at least 4 localizations. We also generally applied filtering by the center-frequency ratio (CFR)[1,4], limiting further analysis to raw MINFLUX localizations with CFR ≤ 0.75.

*Docking strand locations* were obtained from raw MINFLUX localizations by combining all localizations from the same "trace" (TID) to obtain a single localization that represents the location of the docking strand on the sdAB marker. The resulting docking strand locations have a higher localization precision and each such location is calculated as the centroid of the raw MINFLUX localizations belonging to its trace [4,5].

*Subunit locations* (molecule locations). To calculate estimates of unique subunit locations when imaging RyR2s with FP insertions in their subunits (i.e. $RyR2_{D4365\text{-}GFP}$ and $RyR2_{T1365\text{-}PATagRFP}$) we used an approach similar to the procedure used previously to reassign filtered MINFLUX localizations to individual molecules[5]. In this three-step procedure we first combine raw MINFLUX localizations to obtain *docking strand locations* as described above. In a second step a DBSCAN clustering of *docking strand locations* is performed to group *locations* that arise from a potentially repeatedly localized docking strand of an sdAB. The distance ε was chosen to be small enough to avoid incorrectly combining dye locations from sdABs on different subunits. Given that *docking strand locations* also have a localization error, typically < 2nm a DBSCAN ε = 1 nm was adopted. In a final step, DBSCAN clustered *docking strand locations* from the same DBSCAN cluster are combined into a single *subunit location* estimate. The location is calculated as the weighted centroid of the *docking strand locations* in the same DBSCAN cluster. This was implemented via the `MergeClumps` module in PYME which combines localization coordinates weighted by localization error.

*Implementation*. Filtering, grouping and combining of localizations were carried out using a PYME localization data processing pipeline with operations represented by "recipes"[2] that comprise a combination of localization data processing modules to implement operations such as DBSCAN clustering, combining ("merging") localizations, filtering, etc as previously described[2].

*Combining localizations*. When combining $N$ localizations with coordinates $x_i,\ i = 1..N$ and corresponding localization precisions $\sigma_i$, a coordinate $x_c$ of the combined location is calculated as the weighted centroid according to

$$x_c = \sum_{i=1}^{N} w_i\, x_i \quad \text{where}$$

$$w_i = \frac{1}{w}\frac{1}{\sigma_i^2} \quad \text{and}$$

$$w = \sum_{i=1}^{N} \frac{1}{\sigma_i^2}$$

The localization error $\sigma_c$ of the combined localization coordinate $x_c$ is

$$\sigma_c = \sqrt{\frac{1}{w}} \quad \text{and therefore one can also write} \quad w_i = \frac{\sigma_c^2}{\sigma_i^2}\ .$$

If all localizations to be combined have the same localization error $\sigma_i = \sigma_0,\ i = 1..N$, e.g. when combining the raw MINFLUX localizations belonging to the same trace (TID), this simplifies to

$$x_c = \frac{1}{N}\sum_{i=1}^{N} x_i \quad \text{and}$$

$$\sigma_c = \frac{\sigma_0}{\sqrt{N}}$$

This localization combining approach is implemented in the `MergeClump` module of `PYMEVisualize`. The calculations are applied to all 3D coordinate axes, i.e. the $x_i$ and corresponding $\sigma_i$ respectively represent the x, y and z coordinates which are all processed in this way.

#### *Visualising 3D MINFLUX data*

*MINFLUX localizations* were visualised in the `PYMEvisualize` point viewer with localizations shown as OpenGL based Gaussian shaped "pointsprites"[2] and colored according to one of several localization properties including z location, acquisition time, cluster size or cluster ID (a unique number given to each cluster) which results in a Gaussian rendering in 3D. In some 3D visualizations *localizations* were rendered as 3D spheres using the `PYMEvisualize` point viewer.

To determine image resolution we calculated Fourier Ring Correlation (FRC)[6,7] or Fourier Shell Correlation (FSC) measures. To calculate the FRC or FSC values, raw MINFLUX localizations were grouped into two groups of localizations using the "Split by Time Blocks for FRC" functionality of `PYMEVisualize` using a Time Block size of 10. For FRC measurements the two groups of localizations were rendered into two 2D images using the Gaussian rendering method of `PYMEVisualize` and a pixel size of 3 nm. FRC resolution was determined as the 1/7 threshold frequency inverse using the FRC module of PYME-extra. For FSC calculations, the two groups of localizations were rendered into two 3D volume images using the Gaussian rendering method of `PYMEVisualize` and a pixel size of 3 nm. The volume images were saved in MRC format using the Python package `mrcfile` which provides a Python implementation of the MRC2014 file format. The data was uploaded to the EMDB FSC server at https://www.ebi.ac.uk/emdb/validation/fsc which calculated the FSC curve from the provided data. FSC resolution was determined as the 1/7 threshold spatial frequency inverse.

##### *Estimates of effective labeling efficiency from MINFLUX 3D NPC data*

To estimate effective labeling efficiency an approach similar to the strategy introduced by Thevathasan et al.[8] was used, but fully extended to 3D. MINFLUX 3D data of Nup96-mEGFP labelled with anti-GFP sdABs modified with DNA-PAINT docking strands were first rendered to a 2D image using Gaussian rendering in PYMEvisualize (Supplementary Fig. 4 A) and locations of NPCs were detected by a semi-manual approach in Fiji[9] and stored as regions-of-interest (ROIs) in ImageJ ROI format. ROIs were read into the PYMEvisualize[2] SMLM data visualizer using the Python roifile package and utilized to generate a mask and label 3D MINFLUX localisations belonging to detected NPCs with a unique ID. For each NPC the associated set of localisations (identified by a unique ID label) were fit to a 3D double ring template using a maximum-likelihood algorithm similar to an approach described previously[10]. A 3D template was constructed from two rings (nominal diameter 107 nm) spaced 50 nm apart in z and convolved with a Gaussian (sigma = 5 nm) in 3D to provide a smooth template for alignment (Supplementary Fig. 4 B). A small evenly distributed probability was added to account for background localizations[10]. We chose a "smooth" model versus a "detailed" NPC template for robust parameter estimation[10], sufficient for robust estimates of labeled versus unlabeled "segments" of the NPC data sets. Similarly to previous work, for robustness we do not rely on resolving the pairs of nearby Nup96 sites (laterally ~12 nm apart) in the NPC structure but rather test for labeling of each of 8 radial "segments" in the NPC structure that each contain the two nearby Nup96 sites[8], and do this separately for cytoplasmic and nucleoplasmic parts of the NPC. Prior to template fitting the MINFLUX localizations are typically slightly shifted relative to the template and also exhibit a visible tilt angle (Supplementary Fig. 4 B, top row).

The optimization algorithm works on the negative logarithm of the likelihood (NLL) function, i.e. the maximum likelihood is found my minimizing the NLL. The NLL for the given set of localizations and current transformation parameters is obtained by evaluating the NLL template function at each transformed localization coordinate and summing the contribution from all localizations that belong to the NPC[10]. To minimize the NLL, 7 parameters are varied which describe coordinate transformations to line up the MINFLUX data with the template: 3 spatial coordinates to define the NPC center, 2 angles in Euler notation describing the rotation of the NPC central axis and 2 scaling factors to account for small variations in ring diameter and inter-ring spacing, respectively. Minimization is performed with a global optimizer to avoid becoming trapped in local minima[10]. In our implementation we used the `basinhopping` function of the `scipy.optimize` package which performs "basin-hopping", a two-phase method that combines a global stepping algorithm with local minimization at each step[11]. The `basinhopping` function was used with the L-BFGS-B method, i.e. the limited-memory BFGS algorithm that approximates the Broyden–Fletcher–Goldfarb–Shanno algorithm (BFGS), extended with simple bound constraints on variables[12].

The transformed coordinates that minimize the NLL are then used for further determination of labelling of NPC segments in the two rings. An example of aligned localisations is shown in Supplementary Fig. 4 B, bottom row. In addition, the optimal fitting parameters are also used to display the transformed template overlaid with the fitted MINFLUX localisations in PYMEvisualize, see Supplementary Fig. 4 C,D. Notably, the central symmetry axis is also displayed in these overlays and it exhibits visible variation in NPC angles across the field of view.

In the next step, rotational alignment of the aligned NPC data is performed as described for 2D data[8] but in our implementation separately for the cytoplasmic and nucleoplasmic rings as illustrated in Supplementary Fig. 4 E. To determine the “segment boundaries” for an NPC data set we calculate a metric that looks at angles of events and assigns the smallest penalty (zero) at the center of a π/4 (45 degree) segment and increases linearly towards 0.5 at the edges of segments. Contributions from all NPC events are summed up to determine a “penalty” measure. Segment boundaries are rotated in a range of 0 to π /4 and the rotation angle that minimizes the total penalty is the rotation that defines the segment borders for the following determination of “labeled” segments. The number of labeled “segments” in the 8-fold symmetry of the NPC structure is determined using a threshold value. In our analysis, a threshold of 1 localisation was used due to the high signal-to-noise in the data and a threshold analysis that confirmed its suitability (shown in Supplementary Fig. 4 H). Each NPC can exhibit from 0 to 16 labeled segments, as the labeling evaluation is carried out separately for cytoplasmic and nucleoplasmic “rings”.

The data from all NPCs in a dataset evaluated in this way were pooled into a cumulative histogram and fitted to a probabilistic model that predicts the relative number of 0 to 16 labeled segments as a function of the fit parameter $p_{LE}$ which denotes the labeling efficiency. The model is obtained by starting from the binomial probability distribution of having exactly $k$ successes when attempting to label $n$ independent sites with the same labeling probability $p_{LE}$

$$f(k, n, p_{LE}) = \binom{n}{k} p_{LE}^{k} (1 - p_{LE})^{n-k}$$

Similarly to previous work, we test for labeling of each segment in the 8-fold symmetry NPC structure with each segment containing two nearby Nup96 sites[8]. Accordingly, the probability $p_{seg,unlabeled}$ that a segment of the NPC structure is unlabeled is obtained from the binomial probability that both Nup96 sites are not labelled

$$p_{seg,unlabeled} = f(0,2, p_{LE})$$

which gives the probability $p_{seg,labeled}$ that at least one of the sites is labeled as

$$p_{seg,labeled} = 1 - p_{seg,unlabeled} = 1 - f(0,2, p_{LE})$$

From this expression we obtain the probability $p(N, p_{LE})$ that $N$ out of 16 segments are labeled as

$$p(N, p_{LE}) = f\left(N, 16, p_{seg,labeled}\right) = f\left(N, 16,1 - f(0,2, p_{LE})\right)$$

and similarly, the cumulative probability $p_c(N, p_{LE})$ which we show in plots and use as fitting function, as

$$p_c(N, p_{LE}) = \sum_{k=0}^{N} p(k, p_{LE})$$

Processing both experimental and model histograms in cumulative form includes the advantage that the curves have a characteristic sigmoidal shape that shifts to the right with increasing $p_{LE}$, aiding convenient visual inspection of effective labeling performance, see Supplementary Fig. 4 F. The best fit parameter $p_{LE}$ together with its uncertainty $\Delta p_{LE}$ are then provided as estimate of effective labeling efficiency.

Suitability of a threshold for deciding labeling of a segment was tested using background data from a region distal from the immediate vicinity of the nuclear envelope. As shown in Supplementary Fig. 4 G, using a nucleoplasmic region above the level of the NPCs (Background region in Supplementary Fig. 4 Gii) $p_{LE}$ was determined first just using the actual NPC localizations and then also (case 1) only using the background region with the same NPC mask obtained from the actual NPC data and (case 2) displacing all the localisations in the background region into the plane of the NPCs (by reducing the z-coordinates of localizations in the background region by 220 nm to "move" them into the NPC region) and thus determination of $p_{LE}$ in the presence of the additional background. This analysis was carried out for 3 typical datasets that exhibit a $p_{LE}$ of 59.1 % (N=3 datasets from 2 independent realisations, Supplementary Fig. 4 Hi) as reference value. Just evaluating the data from the background region alone (case 1) gives an apparent labeling efficiency $LE_{BG}$ of 3.0 % and, more importantly, in the additional presence of the background (case 2), the mean labeling efficiency only increases from the reference value 59.1 % to 59.3 %, i.e. a difference ΔLE of 0.2% resulting from additional background (N=3 datasets from 2 independent realisations, Supplementary Fig. 4 Hii). This very small increase due to background localizations shows that the NPC data is highly specific and that a threshold of one compound localization (obtained from coalescing a trace of raw MINFLUX localizations as described above) is suitable for determining segment labeling. Overall, it suggests that estimated labeling efficiencies are perturbed by background localizations by one percent at most, most likely significantly less.

The whole procedure is implemented in the `PYMEcs.Analysis.NPC` module of the `PYME-extra` package, specifically in the `LLmaximizerNPC3D`, `NPC3D` and `NPC3DSet` classes. A graphical interface is provided by the `NPCcalcLM` module as a plugin for PYMEVisualize.

##### *RyR2 cluster analysis*

RyR2 cluster analysis was performed using DBSCAN clustering of subunit locations, taking the aspect of MINFLUX data into account that DBSCAN clustering of subunit locations, not RyR2 locations per se, is conducted. A distance eps = 100 nm was used to define functional RyR2 clusters, as 100 nm is often taken as the diffusion distance over which $Ca^{2+}$ release from a local RyR2 has a reasonable probability to trigger release from adjacent RyR2s[13,14]. The distance is thus justified based on the functional definition of a RyR2 cluster as a group of RyR2s that will likely jointly open during the time course of a $Ca^{2+}$ spark.

RyR2 cluster sizes obtained by DBSCAN clustering with eps = 100 nm are reported in two ways, (a) as the number of labeled subunits in a cluster and (b) the number of RyR2s in the corresponding clusters using estimates of labeling efficiency $p_{LE}$ for correction when available (primarily with HEK293 cells expressing RyR2-GFP). We obtain a corrected number *nRyR* of RyR2s in the cluster from the observed number of labeled subunit locations *nSU* in the cluster according to

$$nRyR = \frac{nSU}{4}\frac{1}{p_{LE}}$$

##### *Deviation of cluster shapes from a plane*

To illustrate how much RyR2 clusters in peripheral couplings deviate from a flat geometry we show a typical dataset in Supplementary Fig. 7 from several angles in 3D. In Supplementary Fig. 7 A-F it is apparent that clusters have a somewhat curved shape (akin to "shallow cups"), at the surface membrane. In addition, we show a typical cluster with its corresponding alpha shape[15,16] determined using the MATLAB “alphaShape” functionality. As a measure of the scattering of points, we calculated the standard deviation of z-coordinates (~27 nm) which was typically about 10 times larger than localization errors themselves (2-3 nm), indicating considerable deviation from a “flat” RyR2 arrangement, as shown in Supplementary Fig. 7 F & G.

##### *Generation of simulated data from EM tomography data*

Starting with RyR2 cluster morphologies as determined in ventricular myocytes using EM tomography[17,18] we generated simulated labeled subunit locations assuming variable degree of labeling. Briefly, RyR positions were drawn in Inkscape as squares to match the red squares in Fig. 1 from Asghari et al., 2014[17] and Fig. 2 from Asghari et al., 2020[18]. The coordinates of the resulting squares stored in Inkscape SVG format were read in using Python with the ElementTree XML API as implemented in xml.etree.ElementTree and further processed to define the locations where GFP/tagRFP residues are positioned[19,20] (see *Supplementary Fig. 6*). Based on these subunit label locations simulated MINFLUX localizations were generated by adding localisation errors with a standard deviation of 3.5 nm. Various labelling efficiencies were simulated by randomly selecting a subset of these localizations with the fraction chosen according to the $p_{LE}$ value that was to be simulated.

##### *Assembly of molecular schematics*

For marker size schematics and RyR2 labeling schematics, PDB structures were rendered using the Mol* Viewer[21] and assembled in Inkscape to scale, using structures with PDB IDs 12FX, 6JI8, 5SYF, 1IGT, 3M22, 4OGS, 6I2G.

#### Software

Processing was conducted using the PYME software package[2] with apps including `PYMEVisualize` and `PYMEImage` as well as by directly using PYME API functions in Jupyter notebooks (using Python 3) and the PYME plugin package `PYME-extra` (https://github.com/csoeller/PYME-extra).

#### References


1. Schmidt, R., Weihs, T., Wurm, C. A., Jansen, I., Rehman, J., Sahl, S. J. & Hell, S. W. MINFLUX nanometer-scale 3D imaging and microsecond-range tracking on a common fluorescence microscope. *Nat. Commun.* **12,** 1478 (2021).

2. Marin, Z., Graff, M., Barentine, A. E. S., Soeller, C., Chung, K. K. H., Fuentes, L. A. & Baddeley, D. PYMEVisualize: an open-source tool for exploring 3D super-resolution data. *Nat. Methods* **18,** 582–584 (2021).

3. Gwosch, K. C., Pape, J. K., Balzarotti, F., Hoess, P., Ellenberg, J., Ries, J. & Hell, S. W. MINFLUX nanoscopy delivers 3D multicolor nanometer resolution in cells. *Nat. Methods* **17,** 217–224 (2020).

4. Ostersehlt, L. M., Jans, D. C., Wittek, A., Keller-Findeisen, J., Inamdar, K., Sahl, S. J., Hell, S. W. & Jakobs, S. DNA-PAINT MINFLUX nanoscopy. *Nat. Methods* **19,** 1072–1075 (2022).

5. Pape, J. K., Stephan, T., Balzarotti, F., Büchner, R., Lange, F., Riedel, D., Jakobs, S. & Hell, S. W. Multicolor 3D MINFLUX nanoscopy of mitochondrial MICOS proteins. *Proc. Natl. Acad. Sci.* **117,** 20607–20614 (2020).
6. Nieuwenhuizen, R. P. J., Lidke, K. A., Bates, M., Puig, D. L., Grünwald, D., Stallinga, S. & Rieger, B. Measuring image resolution in optical nanoscopy. *Nat. Methods* **10,** 557–562 (2013).
7. Banterle, N., Bui, K. H., Lemke, E. A. & Beck, M. Fourier ring correlation as a resolution criterion for super-resolution microscopy. *J. Struct. Biol.* **183,** 363–367 (2013).
8. Thevathasan, J. V., Kahnwald, M., Cieśliński, K., Hoess, P., Peneti, S. K., Reitberger, M., Heid, D., Kasuba, K. C., Hoerner, S. J., Li, Y., Wu, Y.-L., Mund, M., Matti, U., Pereira, P. M., Henriques, R., Nijmeijer, B., Kueblbeck, M., Sabinina, V. J., Ellenberg, J. & Ries, J. Nuclear pores as versatile reference standards for quantitative superresolution microscopy. *Nat. Methods* **16,** 1045–1053 (2019).
9. Schindelin, J., Arganda-Carreras, I., Frise, E., Kaynig, V., Longair, M., Pietzsch, T., Preibisch, S., Rueden, C., Saalfeld, S., Schmid, B., Tinevez, J.-Y., White, D. J., Hartenstein, V., Eliceiri, K., Tomancak, P. & Cardona, A. Fiji: an open-source platform for biological-image analysis. *Nat. Methods* **9,** 676–682 (2012).
10. Wu, Y.-L., Hoess, P., Tschanz, A., Matti, U., Mund, M. & Ries, J. Maximum-likelihood model fitting for quantitative analysis of SMLM data. *Nat. Methods* 1–10 (2022). doi:10.1038/s41592-022-01676-z
11. Wales, D. J. & Doye, J. P. K. Global Optimization by Basin-Hopping and the Lowest Energy Structures of Lennard-Jones Clusters Containing up to 110 Atoms. *J. Phys. Chem. A* **101,** 5111–5116 (1997).
12. Zhu, C., Byrd, R. H., Lu, P. & Nocedal, J. Algorithm 778: L-BFGS-B: Fortran subroutines for large-scale bound-constrained optimization. *ACM Trans Math Softw* **23,** 550–560 (1997).
13. Sobie, E. A., Guatimosim, S., Gómez-Viquez, L., Song, L.-S., Hartmann, H., Jafri, M. S. & Lederer, W. J. The Ca2+ leak paradox and rogue ryanodine receptors: SR Ca2+ efflux theory and practice. *Prog. Biophys. Mol. Biol.* **90,** 172–185 (2006).
14. Baddeley, D., Jayasinghe, I. D., Lam, L., Rossberger, S., Cannell, M. B. & Soeller, C. Optical single-channel resolution imaging of the ryanodine receptor distribution in rat cardiac myocytes. *Proc. Natl. Acad. Sci. U. S. A.* **106,** 22275–22280 (2009).
15. Edelsbrunner, H., Kirkpatrick, D. & Seidel, R. On the shape of a set of points in the plane. *IEEE Trans. Inf. Theory* **29,** 551–559 (1983).

16. Edelsbrunner, H. & Mücke, E. P. Three-dimensional alpha shapes. *ACM Trans Graph* **13,** 43–72 (1994).
17. Asghari, P., Scriven, D. R. L., Sanatani, S., Gandhi, S. K., Campbell, A. I. M. & Moore, E. D. W. Nonuniform and variable arrangements of ryanodine receptors within mammalian ventricular couplons. *Circ. Res.* **115,** 252–262 (2014).
18. Asghari, P., Scriven, D. R., Ng, M., Panwar, P., Chou, K. C., van Petegem, F. & Moore, E. D. Cardiac ryanodine receptor distribution is dynamic and changed by auxiliary proteins and post-translational modification. *eLife* **9,** e51602 (2020).
19. Liu, Z., Zhang, J., Li, P., Chen, S. R. W. & Wagenknecht, T. Three-dimensional Reconstruction of the Recombinant Type 2 Ryanodine Receptor and Localization of Its Divergent Region 1*. *J. Biol. Chem.* **277,** 46712–46719 (2002).
20. Liu, Z., Zhang, J., Wang, R., Wayne Chen, S. R. & Wagenknecht, T. Location of Divergent Region 2 on the Three-dimensional Structure of Cardiac Muscle Ryanodine Receptor/Calcium Release Channel. *J. Mol. Biol.* **338,** 533–545 (2004).
21. Sehnal, D., Bittrich, S., Deshpande, M., Svobodová, R., Berka, K., Bazgier, V., Velankar, S., Burley, S. K., Koča, J. & Rose, A. S. Mol* Viewer: modern web app for 3D visualization and analysis of large biomolecular structures. *Nucleic Acids Res.* **49,** W431–W437 (2021).

#### Supplementary Figures

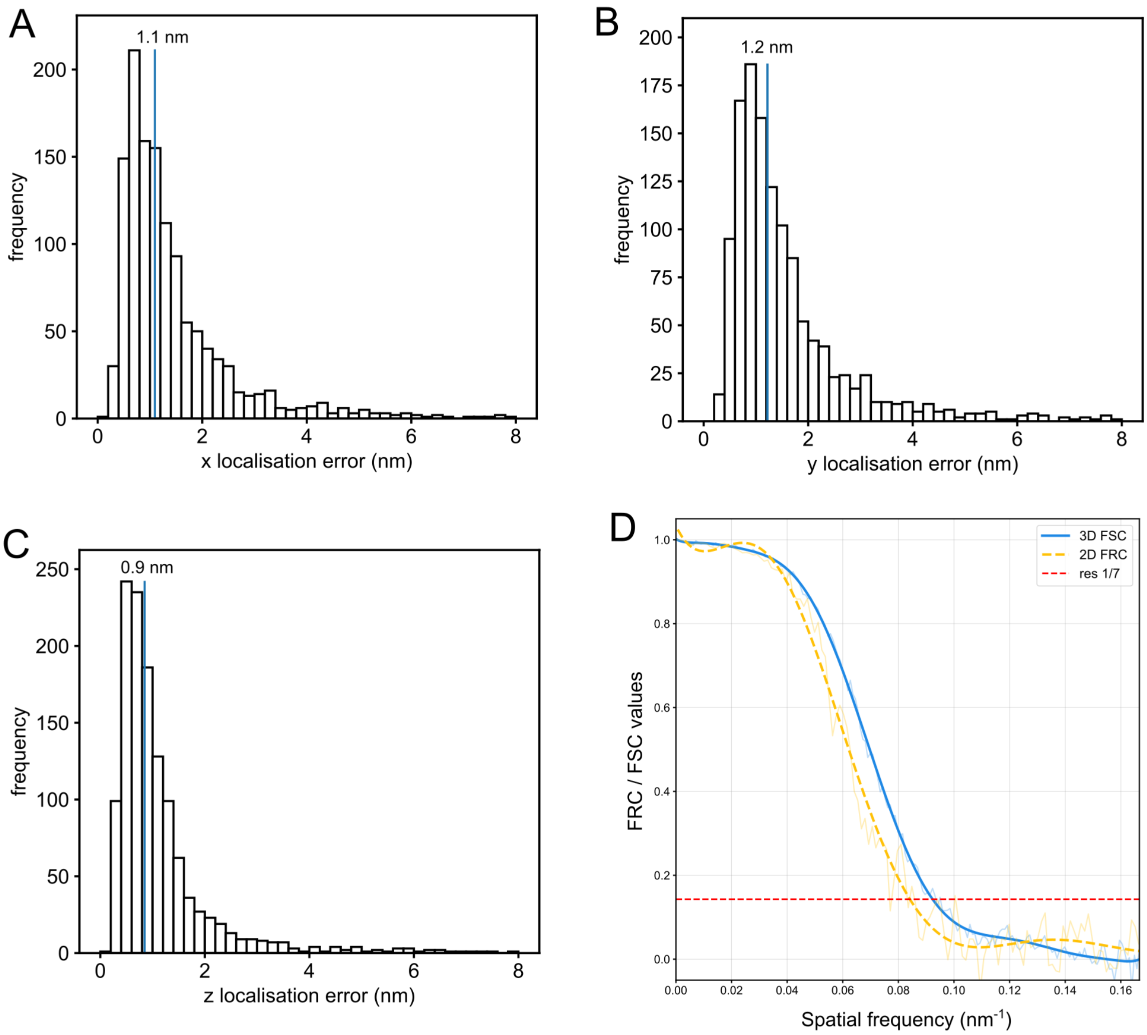


Supplementary Figure 1. Localisation errors and resolution measures. **A-C**. Shown are histograms of x, y and z localization errors of docking strand locations that were obtained by combining raw MINFLUX localization as described in the Methods. Note that median localization errors are well below 2 nm in all directions. **D**. Fourier ring correlation (FRC) and Fourier shell correlation (FSC) of a cardiac myocyte $RyR_{T1365\text{-}PATagRFP}$ MINFLUX dataset, calculated as detailed in the methods. The FRC measurement encompassing only the two-dimensional data yields a 2D resolution of 11.9 nm. The FSC which incorporates the same xy data as the FRC but also includes z coordinates after rendering into a volume data set measures a 3D resolution of 10.8 nm. The similarity between the FRC and FSC resolution values is a further demonstration of the near homogeneous localization of docking strands in all directions when using 3D MINFLUX DNA-PAINT.

A

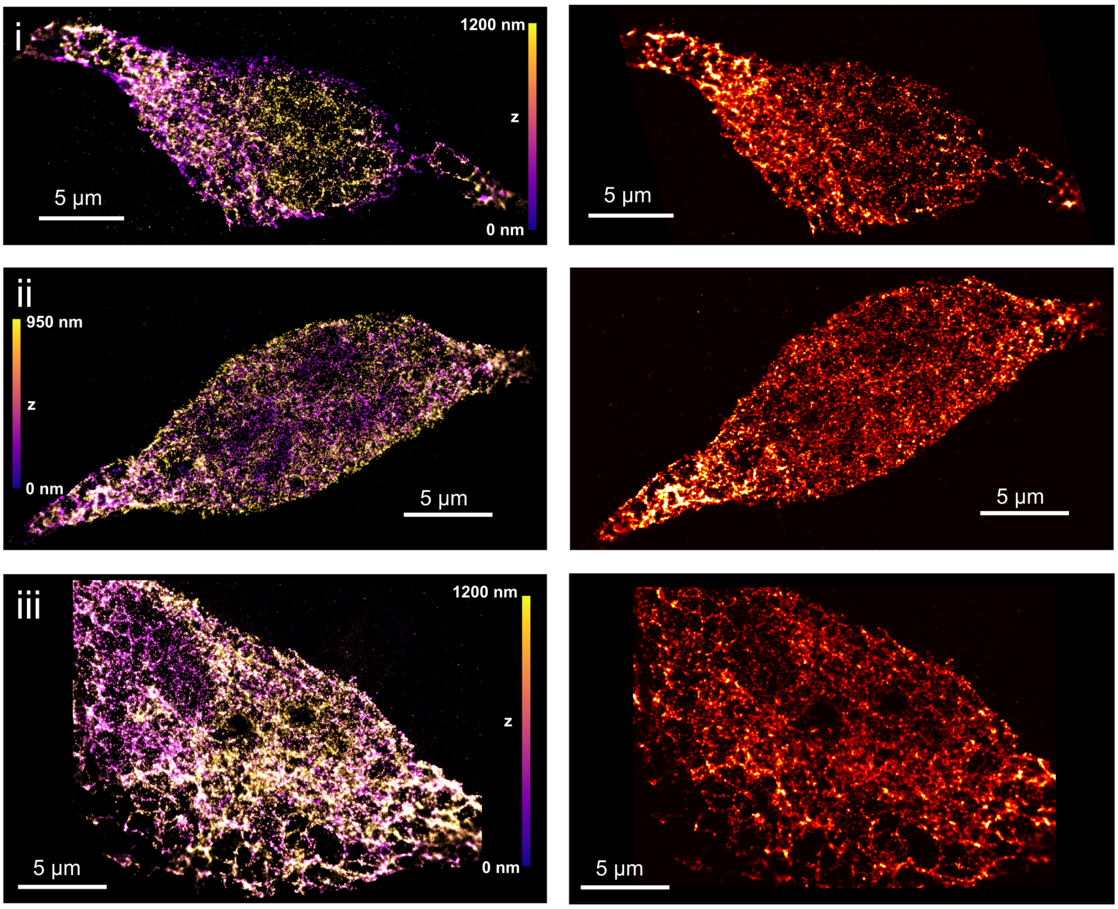


Supplementary Figure 2. Widefield DNA-PAINT data of HEK293 cells stably expressing RyR2$_{D4365\text{-}GFP}$. **A**. Whole cell overviews using 3D widefield DNA-PAINT super-resolution imaging with biplane axial localisation. Left panels show OpenGL based PYMEVisualize 3D renderings with axial position colour-coded, right panels show Gaussian 2D renderings of the corresponding panels on the left.

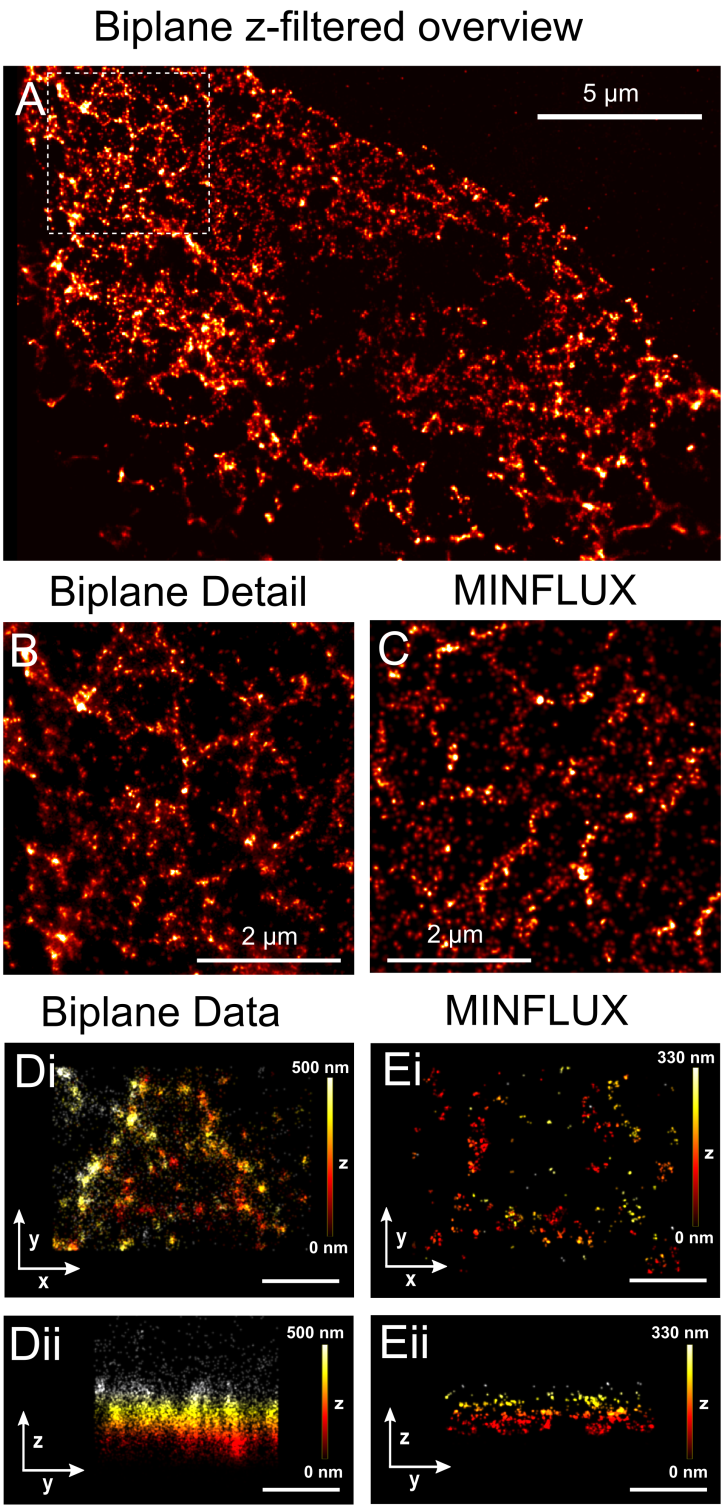


Supplementary Figure 3. Comparison between MINFLUX and wide field DNA-PAINT data. Part 2 – **A**. Overview of the data as in Supplementary Fig. 2 Aiii but filtered to only include localizations in an axial range of 500 nm to match the corresponding MINFLUX dataset. **B**. Magnified detail from B (indicated by box in B). **C**. An area of equivalent lateral extent from an MINFLUX DNA-PAINT dataset, rendered and blurred with a Gaussian of 15 nm diameter (standard deviation) to more closely match the lower resolution in the widefield DNA-PAINT dataset. Note the qualitatively similar appearance. **D** and **E**. Comparison of x-y (i) and x-z (ii) views from widefield biplane data (D) and MINFLUX 3D data (E). Note the much poorer z-detail in the widefield DNA-PAINT data. Scale bars D, E: 500 nm.

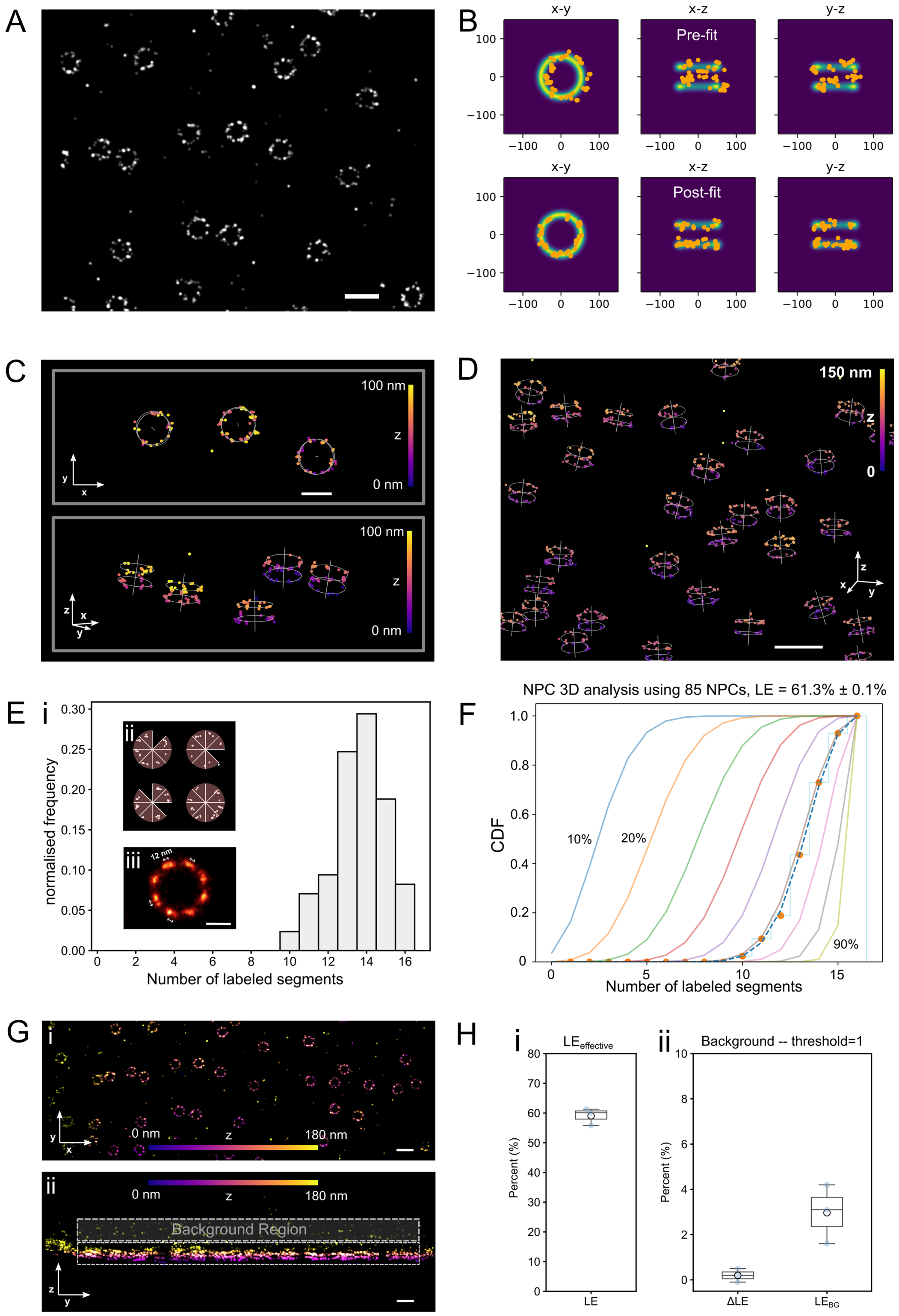


Supplementary Figure 4. Determination of effective labeling efficiency with NPC structures obtained from NPU96-meGFP labeling. **A**. A 2D rendering of the MINFLUX 3D data was rendered and processed in Fiji to detect NPC structures. **B**. Localisations belonging to identified NPC structures were fit using a maximum-likelihood approach implemented with a double ring template.The top row shows various views of the unaligned data from a single NPC prior to fitting and the bottom row indicates the best fit transformation aligning the data with the template. **C**. The best fit data was used to generate template overlays (double ring plus a line indicating the central symmetry axis through the pore). **D**. In this larger field-of-view, otherwise as in C, the variation in NPC axis alignment across the field is visible. **E**. Labeling of segments was determined to generate an experimental histogram (i) following rotational alignment of fitted data to segment boundaries (ii). The aligned NPC localisations can also be overlaid to produce and render an average distribution of Nup96 localisations (iii) which clearly exhibits the 8-fold symmetry and also shows ~12nm distal peaks resulting from the two Nup96 sites in a segment. **F**. Fitting of the cumulative experimental histogramn (dots) to the model yields an estimate of effective labeling efficiency as best fit (dashed line), here 61.3 % ± 0.1 %. **G** and **H**. Testing of labeling threshold against background contribution as described in text, supporting the suitability of a threshold of 1. Scale bars A, D, G: 200nm, C: 100 nm, E: 50 nm,

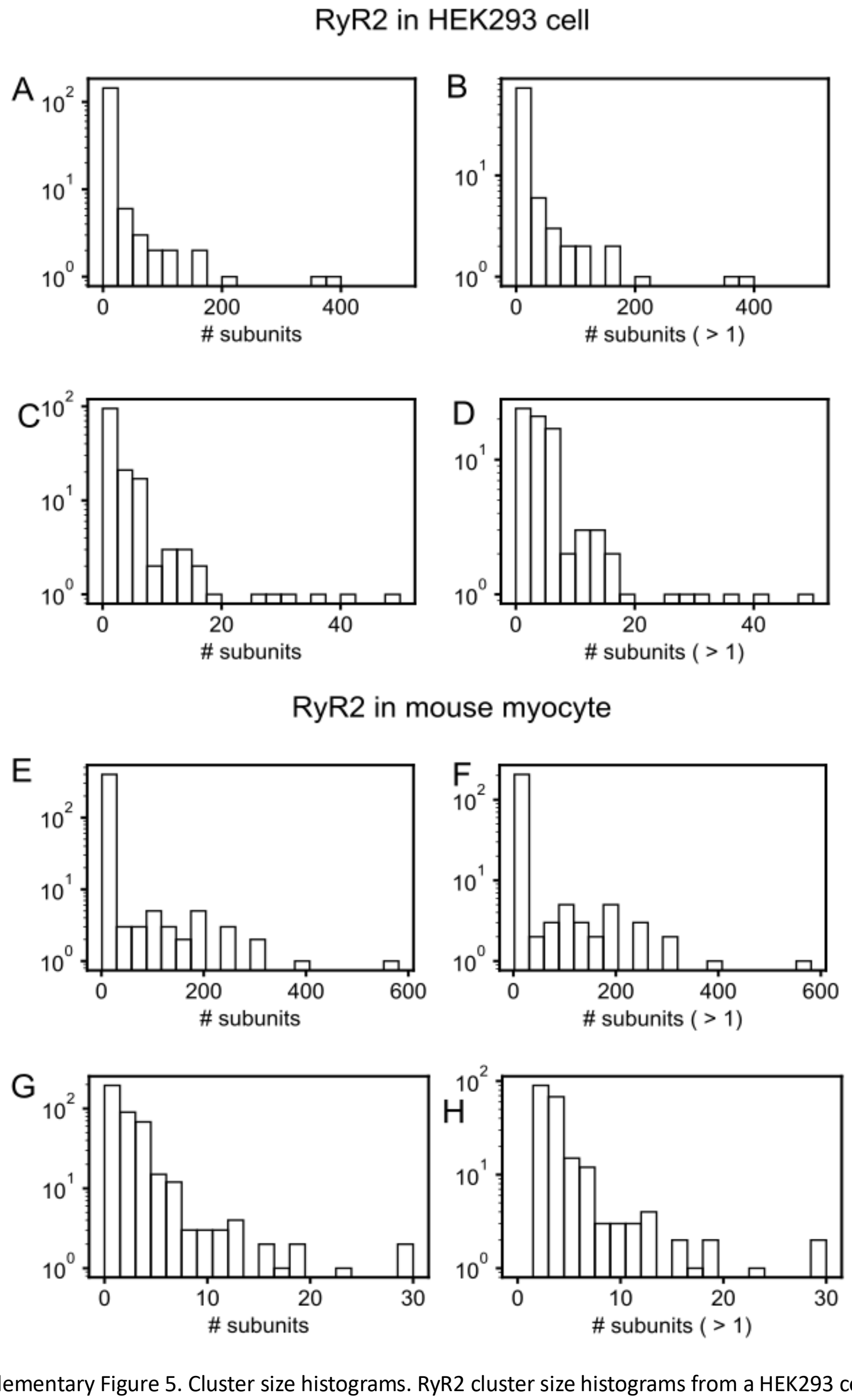


Supplementary Figure 5. Cluster size histograms. RyR2 cluster size histograms from a HEK293 cell (A-D) versus corresponding histograms from a mouse myocyte (E-H). Shown are detected subunit numbers using all clusters (left) and only including clusters with 2 or more subunits (right), highlighting the large number of "isolated subunits", i.e. clusters containing a single or very few subunits. C, D and G, H show the same data as the corresponding panels above, respectively, but focusing on "small" clusters omitting the tail of large clusters that are prominent in visualisations of the data. Note the logarithmic display of cluster numbers.

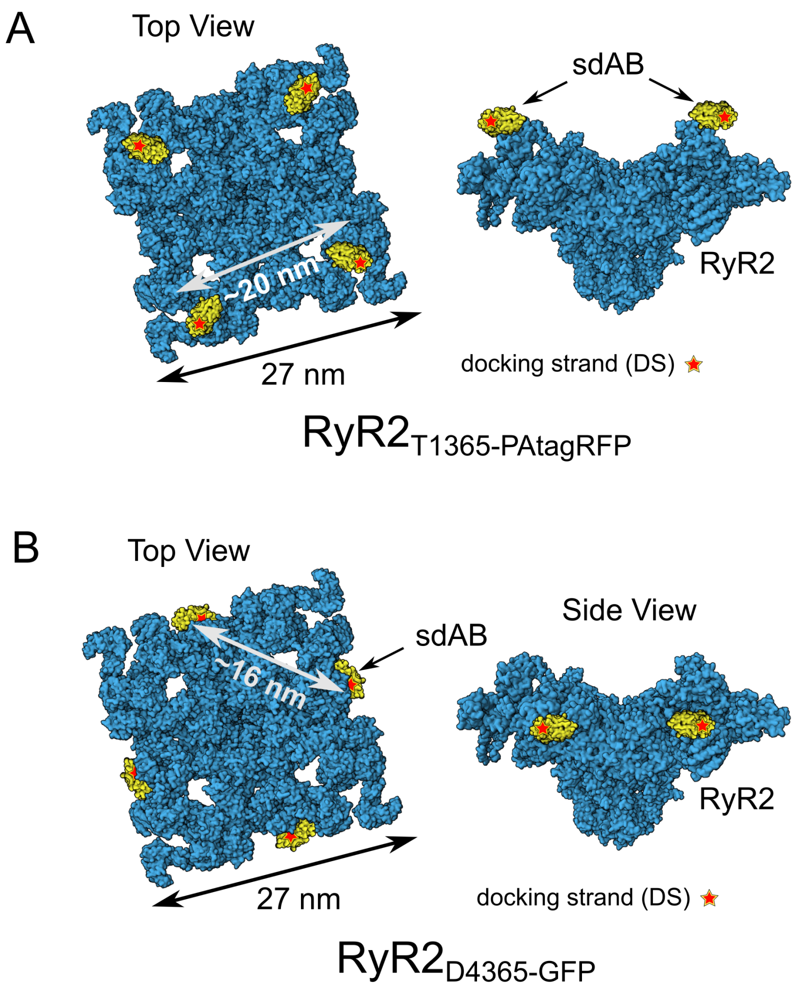


Supplementary Figure 6. RyR2 labeling schematics. **A**. RyR$_{T1365-PATagRFP}$. The location of PA-TagRFP inserts and bound MP anti-TagFP used in PA-RFP RyR2 knock-in mice for both top-down (left) and side views (right). Inserts are situated close to the four corners and are spaced ~20 nm between vertices. **B**. RyR2$_{D4365-GFP}$ . The location of GFP residues on RyR2 tetramers and the corresponding single domain antibody location in HEK293 cells are expressed close to the midpoint of the edges of the RyR2 tetramer with nearest distances between GFP molecules of ~16 nm. Position of sdAB markers are shown based on GFP domain locations read out from figures in references [19] and [20].

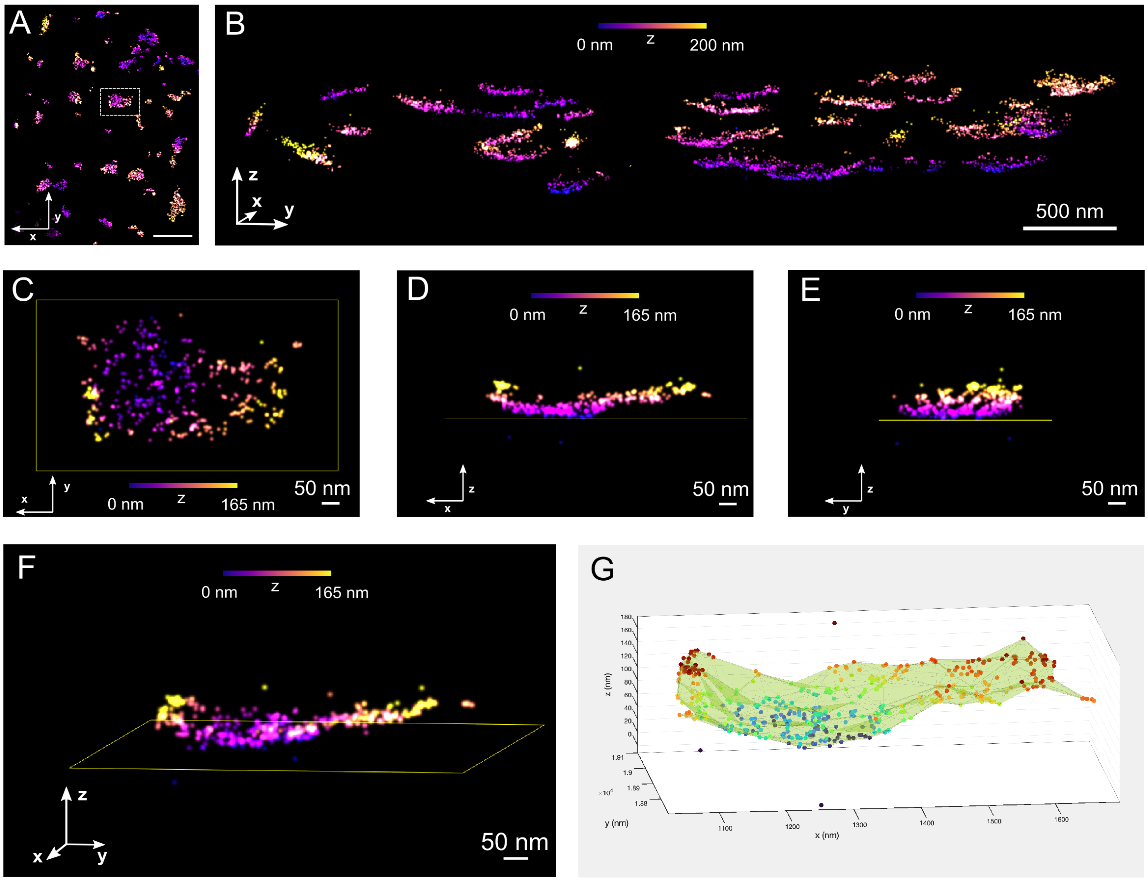


Supplementary Figure 7. Cluster 3D shapes as seen in in 3D views of MINFLUX 3D data in a mouse ventricular myocyte from a tagRFP RyR2 mouse. **A**. X-y overview of a typical dataset. **B**. 3D view of all large clusters (containing >50 subunits). **C.-F**. 3D views of a single cluster exhibiting a shape typically observed, i.e. curving gently away from the coverslip surface. The standard deviation of z locations in this cluster is 27 nm. **G**. Analysis of alpha-shape volume of the cluster shown in C-F, visualized using the MATLAB “alphaShape” functionality with default parameters. Outliers in the z position were detected as values that are more than three scaled median absolute deviations from the median position, using the MATLAB "isoutlier" function. A-G: color indicates z elevation. Scale bar in A: 1 µm.

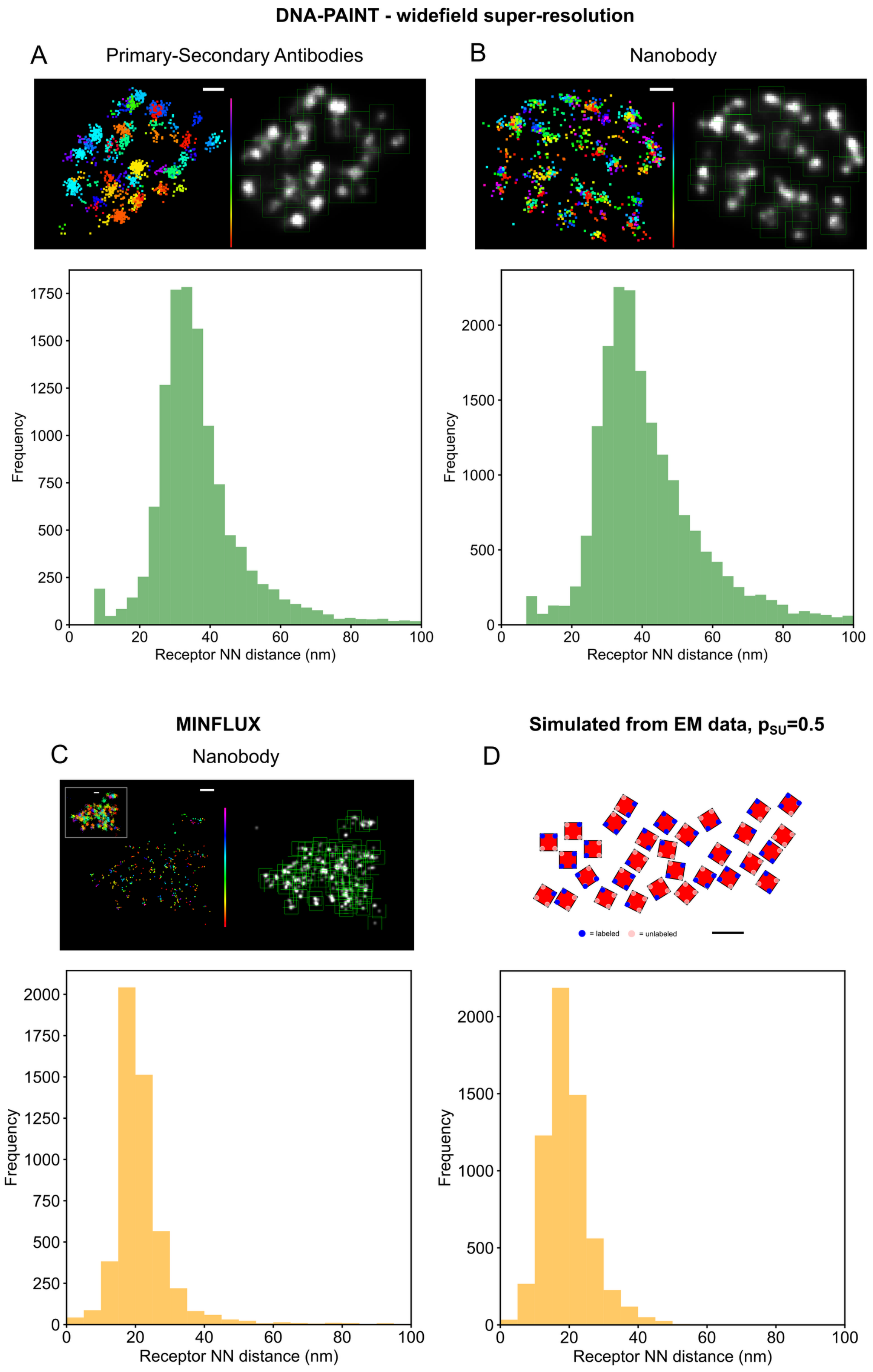


Supplementary Figure 8. Nearest neighbor (NN) distance analysis of RyR2 staining "blobs" in different staining and super-resolution imaging modalities in peripheral couplings of isolated mouse cardiomyocytes. Example localization data of RyR2 clusters are color coded for when in time the events were detected and accompanying rendered images with "blob" detection boxes (green) overlayed, see methods. The resulting nearest neighbor histograms from using blob centroid locations show similar distributions between labeling approaches in widefield super-resolution data but differ between imaging modalities. **A**. DNA-PAINT data for RyR2s labeled with primary & secondary antibodies had a median value of 34.2 nm and mode at 30 nm (N=5 cells from n=2 animals) compared to **B**, a median of 38.1 nm and mode at 35 nm (N=9 cells from n=3 animals) for single domain antibodies, which are not statistically different (p=0.31). **C**. The MINFLUX DNA-PAINT data, which also used the single domain antibody, had a median nearest neighbor distance of 20.0 nm (smaller than widefield DNA-PAINT based imaging measures with ABs or sdABs, p<0.001, respectively) and modal distance of 20 nm (N=5 cells from n=3 animals). **D**. Comparison simulation data generated from EM tomography data, references [17,18], with a labeling efficiency of 0.5 and localization error of 3.5 nm, resulted in a distribution very similar to the MINFLUX data, with median value 18.5 nm and mode at 18 nm. Scale bars A-C: 50 nm, D: 100 nm.
